# Visual awareness judgments are sensitive to accuracy feedback in stimulus discrimination task

**DOI:** 10.1101/572503

**Authors:** Marta Siedlecka, Michał Wereszczyński, Borysław Paulewicz, Michał Wierzchoń

## Abstract

In this study we tested the hypothesis that perceptual awareness judgments are sensitive to the accuracy feedback about previous behaviour. We used a perceptual discrimination task in which participants reported their stimulus awareness. We created two conditions: No-feedback and Feedback (discrimination accuracy feedback was provided at the end of each trial). The results showed that visual awareness judgments are related to the accuracy of current and previous responses. Participants reported lower stimulus awareness for incorrectly versus correctly discriminated stimuli in both conditions; they also reported lower stimulus awareness in trials preceded by incorrect discrimination responses, compared to trials preceded by correct discrimination. This difference was significantly stronger in the Feedback condition. Moreover, in the Feedback condition we also observed larger post-error slowing for PAS ratings. We discuss the relation between the effects of performance monitoring and visual awareness and interpret the results in the context of current theories of consciousness.

## 1.1. Introduction

Can previous mistakes influence our visual awareness? Although the factors that modulate visual perception have been extensively studied, less is known about how visual experience is formed. Studies that investigated subjective aspects of vision have shown that visual awareness of a given stimulus depends not only on sensory data, but also on the perceiver’s attention (e.g. Koivisto & Silvanto, 2011; Koivisto & Silvanto, 2012), expectations (Melloni, Schwiedrzik, Muller, Rodriguez & Singer, 2011; Pinto, van Gaal, de Lange, Lamme, & Seth, 2015), and even stimulus-related actions (Siedlecka, Hobot, Skóra, Paulewicz, Timmermans, & Wierzchoń, 2019). Following recent findings on the relation between motor activity and subjective aspects of perception (Fleming, Maniscalco, Ko, Amendi, Ro, & Lau, 2015; Gajdos, Fleming, Saez Garcia, Weindel, & Davranche, 2019; Siedlecka et al., 2019), we attempted to investigate whether judgments of visual experience are sensitive to internal and external evaluation of one’s performance in perceptual tasks.

Perception processing can be assessed by objective measures, such as forced-choice discrimination or signal detection. Perceptual awareness, however, is typically accessed through first-person reports given by the perceivers. For example, during a perceptual decision task, participants might be asked to rate the clarity of their visual experience using the Perceptual Awareness Scale (PAS, Ramsøy & Overgaard, 2004) or simply report stimulus visibility (Sergent & Dehaene, 2004). Indirect measures such as confidence in stimulus-related choices are also used (Cheesman & Merikle, 1986). In recent years it has been shown that perceptual confidence correlates with some characteristics of a stimuli-related response, such as response time in a discrimination task (Fleming, Massoni, Gajdos, & Vergnaud, 2016; Kiani, Corthell, & Shadlen, 2014) or the occurrence of preparatory muscle activity (Gajdos et al., 2019), and is affected by changes in cortical motor representations (Fleming et al., 2015). Action has also been observed to influence visual awareness ratings (Siedlecka et al., 2019). In this study, participants rated their perceptual awareness of near-threshold stimuli after carrying out a cued response that was either congruent, incongruent or neutral in respect to the correct stimulus-related response. Lower PAS ratings were observed in the neutral condition, thus suggesting that stimulus-related motor response elevates visual awareness of this stimulus.

If awareness of a stimulus can be affected by stimulus-related response, does it also depend on evaluation of the previous action? Such a possibility is allowed by the theories that stress the role of learning in shaping visual awareness. For example, the higher-order Bayesian decision theory of consciousness (Lau, 2008) assumes that the brain learns to distinguish which of its own internal states represent the external stimuli. As a result of this learning process, the criterion for conscious detection might change, thus increasing or decreasing the stimulus awareness level for the same stimulus (Lau and Passingham, 2006). Cleeremans and colleagues (Cleeremans, 2011; Timmermans, Schilbach, Pasquali & Cleeremans, 2012) describe consciousness as a result of the brain learning the neural consequences of its own activity and actions, and increasing the precision of its representations (whether or not it sees something). Learning can therefore improve the ability to access perceptual information; this prediction was supported by a metacontrast masking study in which extensive training in a discrimination task with accuracy feedback increased perceptual awareness for correctly discriminated stimuli (Schwiedrzik, Singer, & Melloni, 2009).

The question asked in this paper is whether perceptual awareness is related to changes in perceptual performance level, specifically whether it is sensitive to the outcome of on-line performance monitoring. Research on motor control indicates the existence of systems that are specialized in monitoring and regulating task-related behaviour and that detect difficulties and errors in order to adjust the level of top-down control (Botvinick, Cohen, & Carter, 2004; Ridderinkhof, Ullsperger, Crone, & Nieuwenhuis, 2004). The results of monitoring are not necessarily consciously perceived (Endrass, Reuter, & Kathmann, 2007; Nieuwenhuis, Schweizer, Mars, Botvinick, & Hajcak, 2007; Wessel, Danielmeier, & Ullsperger, 2011), but they might affect subsequent behaviour. For example, after an error is committed in speeded response tasks, participants respond more slowly (so-called post-error slowing, Dutilh, Vandekerckhove, Forstmann, Keuleers, Brysbaert, & Wagenmakers, 2012; Notebaert, Houtman, Van Opstal, Gevers, Fias, & Verguts, 2009; Rabbitt, 1966) and make fewer mistakes, therefore they adapt their performance speed in order to achieve a certain level of accuracy (Veen & Carter, 2006). There are several theories explaining how errors can be detected, but the most popular ones state that it stems from the comparison between the required and the given response (Coles, Scheffers, & Holroyd, 2001; Falkenstein, Hoormann, Christ, & Hohnsbein, 2000; Gehring, Gross, Coles, Meyer, & Donchin, 1993) or from the conflict between simultaneously evolving response tendencies (Botvinick, Braver, Barch, Carter, & Cohen, 2001; Botvinick, et al., 2004). Erroneous responses are accompanied by electrophysiological changes in brain activity, such as error-related negativity (ERN, Gehring et al., 1993) followed by a positive component, Pe (see e.g. Boldt & Yeung, 2015; Wessel, 2012). Similarly, explicit information about an error is accompanied by feedback-related negativity (FRN, e.g. Luu, Tucker, Derryberry, Reed, & Poulsen, 2003) and has also been shown to be related to subsequent behaviour (Derryberry, 1991; Luu et al., 2003).

It is not clear what the relation is between the results of performance monitoring and awareness. Studies on confidence in perceptual decisions show that confidence level is related to decision accuracy (Kiani et al., 2014; Petrusic & Baranski, 2003), but this relationship could simply reflect the influence of sensory data quality on both objective and subjective aspects of perception (the less perceptual evidence there is, the higher the probability of making mistakes and the lower the level of stimulus awareness). Also, studies on error detection are not consistent when it comes to the relation between error-related neural activity and error awareness, although the amplitude of ERN and Pe has been shown to correlate with choice confidence and error detection (Bold & Yeung, 2015; Scheffers & Coles, 2000; Steinhauser & Yeung, 2012). However, it has been proposed that information about errors (conscious or unconscious) and other action-related information could be integrated with data from the visual system in the prefrontal cortex, thus resulting in decreased stimulus awareness (Anzulewicz, Hobot, Siedlecka, & Wierzchoń, 2019).

In this study we tested the hypothesis that visual awareness judgments are related to the results of performance monitoring, and specifically to internal or external accuracy feedback about previous behaviour. We used a perceptual discrimination task in which participants also reported their stimulus awareness. Half of the trials were followed by explicit feedback about discrimination accuracy. We expected that reported awareness level would be related to the accuracy of the current and previous trials. Specifically, we hypothesized that awareness ratings would decrease after erroneous trials, especially when participants received accuracy feedback. However, it is also plausible that committing an error or receiving negative feedback raises the reported awareness level by increasing top-down attention and amplifying the related sensory signal (Fazekas & Overgaard, 2018).

## 2.1. Methods

### 2.2. Participants

Thirty-seven healthy volunteers (7 males) aged 22 (SD = 2.88) took part in the experiment in return for a small payment. All participants had normal or corrected-to-normal vision and gave written consent to participation in the study. The ethical committee of Jagiellonian University Institute of Psychology approved the experimental protocol.

### 2.3. Materials

The experiment was run on PC computers using PsychoPy software (Peirce, 2007). We used LCD monitors (1280 × 800 pixels resolution, 60Hz refresh rate). The keyboard buttons were labelled for orientation responses (“L” and “R” on the left side of the keyboard) and PAS scale responses (numbers “1”–“4” on the right side of the keyboard). The stimuli were Gabor gratings embodied in visual noise and oriented towards the left (−45 degrees) or the right (45 degrees), presented in the centre of the screen against a grey background. The visual angle of the stimuli was around 3°. The contrast of the stimuli was determined for each participant during a calibration session.

The PAS was presented with the question ‘How clear was your experience of the stimulus?’; the options were ‘no experience’, ‘a brief glimpse’, ‘an almost clear experience’, and ‘a clear experience’. The meaning of the individual scale points was explained in the instruction. The description of each point was based on a guide by Sandberg & Overgaard (2015) with some modifications related to the characteristics of stimuli that were relevant in this experiment (i.e. “no experience” was associated with no experience of the Gabor stripes, but “a brief glimpse” was associated with an experience of “something being there” but without the ability to determine the orientation of the stripes).

### 2.4. Procedure

The experiment was run in a computer laboratory. Two within-subject conditions were introduced: with and without accuracy feedback. All trials began with a blank presentation (500 ms), followed by a fixation cross (500 ms). The grating embedded in white noise was presented for 33 ms. Participants were asked to decide whether the grating was oriented towards the left or the right side. The first part of the experiment started with 15 training trials in which the stimuli was clearly visible (presented in colour in RGB space = [0.3,0.3,0.3] and opacity = 1). Participants were asked to discriminate Gabor orientation and they received accuracy feedback. Subsequently, the calibration procedure was used to estimate the stimulus contrast that resulted in about 70% of discrimination responses being correct. There were 150 trials with a 1-up, 2-down staircase (stair-size 0.005, limit for 0.02 and 0.08) and the contrast was established based on the last 100 trials. The calibration procedure was followed by a break. Afterwards, the second session started with 15 training trials in which the PAS scale was presented after the discrimination response.

After the second training session, two experimental blocks followed: with feedback and without feedback (the order was counterbalanced between participants). Each experimental block consisted of 200 trials. Participants were asked to respond to the discrimination task and then to PAS. In the feedback condition, information about discrimination accuracy was presented after the PAS rating. The time limit for all responses was 3 seconds. Participants were asked to respond as quickly and as accurately as possible. The duration of feedback presentation was half a second. The outline of the procedure is presented in Figure 1.

**Figure 1.**
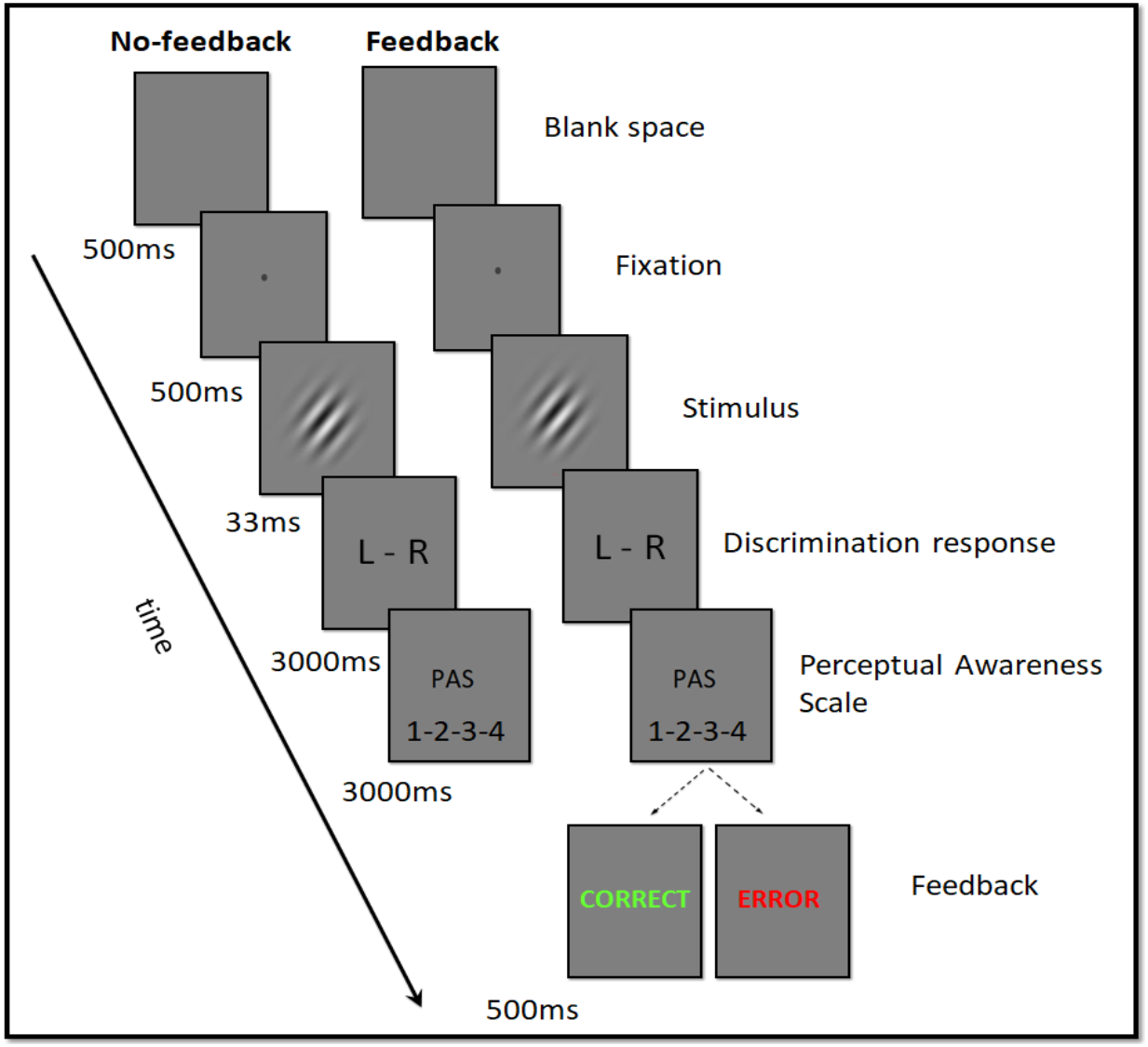
The outline of the procedure. Participants started with either the Feedback or No-feedback block.

## 3.1. Results

We excluded 1 participant’s data from analysis due to poor performance (49% accuracy). Next, we removed omitted trials (discrimination and PAS rating) and discrimination response times shorter than 100 ms. We did not detect significant differences between conditions in accuracy (Feedback, 78%; No-feedback, 75%; *z* = −1.62, *p* = .11). Signal-detection analyses conducted on the discrimination task did not reveal statistically significant differences between conditions in d’ (*z* = −1.31, *p* = .19) or in response bias (*z* = 1.48, *p* = .14).

### 3.2. Confirmatory analyses

#### 3.2.1. The relation between current trial accuracy and PAS ratings

To test the relationship between the accuracy of a given trial and PAS ratings, we fitted a mixed logistic regression model using the lme4 package in the R Statistical Environment (Bates, Maechler, Bolker, & Walker, 2015; R Core Team, 2015). Statistical significance was assessed with the Wald test. The model included the following fixed effects: PAS rating, condition, their interactions, as well as the random effects associated with all the fixed effects. The intercept informs about performance level in No-feedback condition when participants report having no experience of the stimulus (criterion), while the regression slope reflects the relation between PAS rating and discrimination accuracy. The results show a statistically significant relationship between accuracy and PAS ratings, meaning that correct responses were followed by higher PAS ratings in No-feedback condition (*z* = 6.58, *p* < .001); we did not detect significant differences between the conditions in the strength of this relation (*z* = 0.31, *p* = .76). Additionally, we observed that participants’ performance level in No-feedback condition was significantly higher than chance level when they reported having no experience of the stimulus (*z* = 3.36, *p* < .001), but we have not found significant differences in this respect between conditions (*z* = 0.14, *p* = .89). The results are presented in the Table 1. We also performed another analysis that included only the first blocks in each condition because during the Feedback block participants had a chance to improve their metacognitive awareness before getting to the No-feedback block. However, this analysis revealed the same pattern of results.

**Table 1.** The effect of PAS and condition on trial accuracy: mixed logistic regression model summary.

| Number of participants: 36<br>Number of observations: 14,248 | Estimate | SE | $z$ | $p$ |
| --- | --- | --- | --- | --- |
| **Intercept** | 0.37 | 0.11 | 3.36 | < .001*** |
| **PAS rating** | 2.33 | 0.35 | 6.58 | < .001*** |
| **Feedback** | 0.02 | 0.12 | 0.14 | 0.89 |
| **PAS rating: Feedback** | 0.10 | 0.32 | 0.31 | 0.76 |

#### 3.2.2. The relation between previous trial accuracy and PAS ratings

This analysis was carried out to determine whether stimulus awareness ratings are related to the previous trial accuracy and whether this relationship depends on the occurrence of feedback. We compared PAS ratings in post-correct trials (current trials following correct trials) to PAS ratings in post-error trials (current trials following erroneous trials). Linear mixed model with random participant-specific effects of condition and intercept (with effect of previous accuracy nested within conditions, see Table 2) showed that in both conditions participants reported lower stimulus awareness in post-error trials (Feedback: *t*(54.71) = 6.38, *p* < .001; No-Feedback: *t*(51.57) = 3.28., *p* < .001). However, a separate model showed this difference was bigger in the Feedback condition (*t*(1029) = 3.1, *p* = .002).

**Table 2.** The effect of previous trial accuracy and condition on PAS ratings: mixed linear regression model summary.

|  |  |  |  |  |  |
| --- | --- | --- | --- | --- | --- |
| Number of participants: 36<br>Number of observations: 14,212 | **Estimate** | **SE** | **df** | **t** | **p** |
| **Feedback** | 0.41 | 0.03 | 35.78 | 11.89 | < .001*** |
| **No-feedback** | 0.39 | 0.03 | 35.32 | 15.12 | < .001*** |
| **Feedback: previous accuracy** | 0.06 | 0.01 | 54.71 | 6.38 | < .001*** |
| **No feedback: previous accuracy** | 0.03 | 0.01 | 51.57 | 3.28 | .002** |

### 3.3. Exploratory analyses

#### 3.3.1. The relation between previous trial accuracy and response times

A common way of measuring post-error slowing is to compare RTs in correct post-correct and correct post-error trials. However, we see several issues with controlling the current trial accuracy in this approach. This is not the place to elaborate on them, but, for example, if current trial accuracy is affected by both previous trial accuracy and by awareness of current stimulus, controlling current trial accuracy might result in a spurious correlation between previous trial accuracy and current PAS (“collider bias”, see i.e. Paulewicz, Siedlecka, & Koculak, 2019). Therefore, to test whether feedback modulated the way in which the accuracy of the previous response related to the discrimination response time in a current trial we included all current trials into the analysis. We fitted a linear mixed model with reaction time as the dependent variable, fixed effects of previous accuracy, condition and their interaction, as well as random effects associated with all fixed effects. We did not observed longer response times in trials preceded by errors, neither in the Feedback condition (*t*(58.17) = −1.59, *p* = .12), nor in the No-feedback condition (*t*(54.66) = 0.14, *p* = 0.9.

To determine whether the latency of the PAS rating was also related to decision accuracy in the previous trial, we fitted a linear mixed model with PAS response time as dependent variable, fixed effects of previous accuracy, condition and their interaction, as well as and random effects associated with all fixed effects. The model showed that PAS ratings were slower in post-error trials compared to post-correct trials in the Feedback condition (*t*(>100) = 5.92, *p* < .001). This difference was larger than in the No-feedback condition (*t*(>100) = −4.19, *p* < .001).

## 4.1. Discussion

The question of whether the results of performance monitoring could affect perceptual awareness arises from findings that revealed links between stimulus-related action and subjective reports (Gajdos et al., 2019; Fleming et al., 2015; Kiani et al., 2014; Siedlecka et al., 2019). The presented study shows that visual awareness judgments are related to the accuracy of the current and the previous response. Participants reported lower stimulus awareness for incorrectly versus correctly discriminated stimuli in both conditions, with and without explicit accuracy feedback. Moreover, participants reported lower stimulus awareness in trials preceded by trials in which discrimination was incorrect, compared to trials preceded by correct discrimination. This difference was significantly larger in the condition with accuracy feedback. Moreover, in the Feedback condition we observed post-error slowing for PAS rating: responses were longer for correct trials preceded by an error compared to trials preceded by correct responses.

The results suggest that visual awareness judgements are sensitive to the evaluation of one’s previous performance. Although PAS ratings were slower after errors, thus indicating top-down regulation, the results do not support the alternative hypothesis that increased attention after errors or negative feedback leads to higher stimulus awareness. Rather, it seems that performance evaluation could lead to adjustment of subjective criterion for detection (Lau, 2008) or that information about errors is integrated into a higher-order representation (Cleeremans, 2011; Timmermans et al., 2012). If we adopt the hierarchical view on consciousness, which assumes that an awareness report is a type of metacognitive judgment, we can interpret the results in the context of studies on perceptual confidence which show that such judgments are informed by different types of information (Fleming et al., 2015; Gajdos et al., 2019; Kiani et al., 2014; Siedlecka et al., 2019). For example, Fleming’s model of self-evaluation judgments assumes that metacognitive processes assess not only the internal evidence for a decision, but also one’s performance (e.g. by detecting errors, Fleming & Daw, 2017). Studies on performance monitoring show that errors are not only correlated with changes in electrical brain activity (Boldt & Yeung, 2015; Gehring et al., 1993; Wessel, 2012; Wiens et al., 2011), but also with specific responses of the autonomic nervous system, such as changes in heart rate (Fiehler, Ullsperger, Grigutsch, & von Cramon, 2004; Hajcak, McDonald, & Simons, 2003; Wiens et al., 2011), skin conductance, and pupil diameter (OʼConnell, Dockree, Bellgrove, Kelly, Hester, Garavan, Robertson, & Foxe, 2007; Critchley, Tang, Glaser, Butterworth, & Dolan, 2005). We propose that results of performance monitoring that are gathered from different sources (action preparation, characteristics of motor response, proprioceptive and interoceptive feedback) are integrated into a conscious representation of a stimulus. Studies also show that error-related evidence accumulates over time (Ullsperger, Harsay, Wessel, & Ridderinkhof, 2010), therefore it could be reflected not only in the immediate awareness rating but also in consecutive ones.

In our study, PAS ratings were related to the accuracy of current and previous trials independently of external feedback, but they were also affected by feedback. Not only did feedback increase the difference between PAS ratings in trials following correct and incorrect trials, it also affected the latency of PAS rating. These results suggest that explicit accuracy information adds to the internally generated effects of performance monitoring and results in the speed of response being adjusted (more cautious strategy). However, more studies are needed to more precisely establish the links between accuracy feedback and awareness judgments. Different types of feedback might differentially affect stimuli awareness. For example, in the only study known to us in which the effect of feedback on perceptual awareness was tested, participants first underwent extensive perceptual orientation training (five days) with accuracy feedback but without PAS ratings. In the post-training session, the perceptual sensitivity and stimulus awareness of correctly discriminated stimuli were shown to increase (Schwiedrzik et al., 2009). One could assume that since performance feedback has been shown to increase perceptual accuracy (Herzog, & Fahle, 1997; Seitz, Nanez, Holloway, Tsushima, & Watanabe, 2006; Schwiedrzik et al., 2009), it would also increase awareness. Interestingly, in other studies in which perceptual awareness was not measured, false positive feedback (after the whole block) also improved perceptual discrimination accuracy (Shibata, Yamagishi, Ishii, & Kawato, 2009; Zacharopoulos, Binetti, Walsh, & Kanai, 2014). A promising direction for future studies would be to test the effect of trial-by-trial false feedback in order to dissociate the effects of real discrimination accuracy and feedback information on stimulus awareness ratings.

The study has its limitations. Errors, post-error perceptual awareness level, and PAS response latencies, all of which are just observed but not manipulated, might be correlated due to global fluctuations in performance during the task (Dutilh, Vandekerckhove, Forstmann, Keuleers, Brysbaert, & Wagenmakers, 2012). For example, a participant who loses concentration might become erratic in a few consecutive trials and at the same time experience a lower level of stimulus awareness, thus responding slower. One of the solutions to deal with this confound would be to assess the differences between each correct post-error trial and a correct pre-error trial (which follows another correct trial); however, this method requires that each participant has a large number of trials of each type to obtain statistical power. A solution we used is to manipulate the occurrence of accuracy feedback. If the errors had been due to decreasing attention or motivation, negative feedback should have resulted in improved performance and awareness in following trials. The results suggest the contrary: the relations between the accuracy the previous trial, PAS level, and PAS rating latency were stronger or only detected in the Feedback condition.

Another limitation of the current study is that we did not differentiate between the two types of errors: premature responses, when participants pressed the wrong key by accident, and errors resulting from “data limitation” (Scheffer & Coles, 2000). The first type is detected as an error by a subject, whereas the second type results from uncertainty about the stimulus’s characteristics. We presume that as the time available for responses in our task was quite long (3s) and stimulus duration was short (33 ms), most errors were of the second type, although in the feedback condition participants received clear error information despite the type of error. We cannot exclude the possibility that perceived and non-perceived errors are differently related to the level of stimulus awareness.

Summing up, the results of the present experiment suggest that visual awareness judgments are sensitive to the effects of explicit and implicit accuracy monitoring. Although explanations have been proposed of how and when people become aware of committing an error (e.g. Charles et al., 2013; Wiens et al., 2011), little is known about how and why stimulus awareness might be related to previous erroneous responses. Nevertheless, the results could be seen as challenging the claim that perceptual awareness ratings refer directly to one’s visual experience of a stimulus and not to evaluation of one’s performance (Sandberg, Timmermans, Overgaard, & Cleeremans, 2010).

